# Structural and mechanistic basis of ubiquitous bacterial kinase signaling identifies PorX as a noncanonical substrate in *Porphyromonas gingivalis*

**DOI:** 10.64898/2026.01.15.699779

**Authors:** Anshu Saran, Natalie Zeytuni

**Affiliations:** The Department of Anatomy and Cell Biology, McGill University, Montreal, Quebec, Canada. Address: 3640 Rue University, Montreal, QC, Canada H3A 0C7; Centre de Recherche en Biologie Structurale (CRBS), Montreal, Quebec, Canada

**Keywords:** Ubiquitous bacterial Kinase (UbK), Ubk1, autophosphorylation, transphosphorylation, PorX, response regulator, Type IX Secretion System (T9SS), *Porphyromonas gingivalis*

## Abstract

Protein phosphorylation enables bacteria to coordinate regulatory networks underlying virulence and environmental adaptation. The ubiquitous bacterial kinase family comprises atypical kinases with dual serine threonine and tyrosine specificity, yet their structural organization, catalytic mechanisms, and physiological roles remain incompletely defined. In the anaerobic oral pathogen *Porphyromonas gingivalis*, the sole UbK homolog, UbK1, was previously shown to phosphorylate the orphan response regulator RprY, linking UbK1 to virulence-associated pathways. Here, we present the crystal structure of UbK1, revealing the conserved Walker A, HxDxYR, SPT/S, and EW motifs arranged around the ATP-binding site. Structure-guided mutagenesis establishes essential roles for these motifs in ATP hydrolysis and kinase activity. Phosphosite mapping identifies multiple autophosphorylation sites, with the flexible SPT/S loop showing the strongest occupancy and supporting a model in which loop centered autophosphorylation is a major feature of UbK1 cycling while additional sites arise through intermolecular phosphotransfer in *trans*. Consistent with this, biochemical assays demonstrate that UbK1 undergoes autophosphorylation both in *cis* and in *trans*, arguing against a strictly intramolecular autokinase mechanism. Using conserved gene neighborhood analysis, we identified the orphan response regulator PorX as a previously unrecognized UbK1 substrate, consistent with the relaxed substrate specificity reported for UbK homologs. UbK1 catalyzes PorX transphosphorylation *in vitro* at a single tyrosine residue within the receiver domain, independent of PorX oligomeric state. *In vivo*, *ubk1* deletion causes a modest reduction in gingipain secretion, whereas mutation of the UbK1 dependent PorX phosphosite does not measurably affect T9SS mediated cargo export, indicating that UbK1-PorX signaling is separable from PorX canonical gingipain secretion output. Together, these findings establish a structural and mechanistic framework for UbK1 function, expand the UbK substrate repertoire, and support a model in which UbK1 contributes to regulatory pathways in *P. gingivalis* that extend beyond canonical secretion associated outputs.

## Introduction

*Porphyromonas gingivalis* is a Gram-negative, asaccharolytic obligate anaerobe and a keystone pathogen in the severe inflammatory gum disease chronic periodontitis (1). *P. gingivalis* deploys a wide array of virulence factors that enable it to colonize, integrate into and persist within the polymicrobial biofilms in the subgingival pockets (2). The production, processing, and activity of these factors are governed by complex regulatory networks, many of which rely on post-translational modifications, including phosphorylation of protein effectors and regulatory proteins (3,4). Such phosphorylation events, mediated by protein kinases, provide rapid and reversible mechanisms of signal transduction that allow bacteria to adapt to changing environmental and community contexts (5,6).

In bacteria, protein phosphorylation is mediated by multiple kinase families. Two-component histidine kinases drive the most prevalent phosphorelay pathways, (7) while additional kinase classes support serine, threonine, and tyrosine phosphorylation in diverse regulatory contexts (6,8,9). In *P. gingivalis*, kinase-dependent signaling has been linked to major virulence-related behaviors including heme acquisition, biofilm interactions, and regulation of secreted effectors (4). Beyond these established classes, the Ubiquitous bacterial kinase (UbK) family represents a more recently recognized group of atypical kinases with dual specificity for serine/threonine and tyrosine residues (6). Although UbK kinases are broadly distributed across bacterial genomes, they remain under-characterized. While emerging work links UbK homologs to cellular physiology, including stress adaptation and control of translation-associated processes, (6,10) only a limited number of UbK family members (<6) have been experimentally characterized to date, leaving important questions regarding their structural features, catalytic mechanisms, and physiological roles unresolved (6,10–12). UbK proteins were initially annotated as members of a broadly conserved ATPase family (UPF0079/YdiB–YjeE), but biochemical and structural work established that they function as protein kinases with a unique ATP-binding fold distinct from classical Hanks-type and BY kinases (6). In *Bacillus subtilis*, the prototypic UbK homolog YdiB autophosphorylates and transphosphorylates protein substrates *in vitro*, and genetic and biochemical evidence further links this kinase family to cellular stress tolerance and to protein translation through phosphorylation of the tRNA-modification factor YdiE (6).

UbK proteins are generally small, single-domain kinases that contain four conserved signature motifs, Walker A, HxDxYR, SPT/S and EW (6). Biochemical and mutational analyses of UbK homologs indicate that these motifs cooperate to support ATP binding, autophosphorylation and phosphotransfer. The Walker A motif mediates nucleotide binding, while the HxDxYR and EW contribute to the catalytic architecture required for phosphoryl transfer (6). The SPT/S motif, together with residues within one to two positions downstream, has been linked to autophosphorylation across UbK family members and may also contribute to substrate transphosphorylation (6,11–14).

A recent study identified a single UbK homolog in the *P. gingivalis* genome (13). This protein, Ubk1 (∼15 kDa), was shown to autophosphorylate and subsequently transphosphorylate the orphan response regulator RprY on a tyrosine residue within its receiver domain, a modification linked to regulation of heterotypic community development and virulence in *P. gingivalis* (15) (16). Notably, UbK1 was reported to be essential in *P. gingivalis* strain ATCC 33277, as attempt to generate deletion strain in this background were unsuccessful (13). Despite this initial characterization, several important gaps remain in our understanding of UbK1 function. First, the absence of structural information limits the mechanistic insight into how the conserved signature motifs contribute to phosphorylation events, particularly for reported phosphosites predicted to lie far from the catalytic pocket. Second, the UbK1 interactome remains poorly defined, with only a single substrate identified to date. Given parallels with BY kinases and previous studies on UbK homologs, (4,6) it is plausible that UbK1 also exhibits relaxed substrate specificity and engages additional protein partners beyond RprY. Identifying such partners is essential for understanding how UbK1 is integrated into broader regulatory signalling networks in *P. gingivalis*.

Based on these considerations, we hypothesized that UbK1 signaling might intersect with pathways involving the orphan response regulator PorX. In particular, conserved gene neighborhood organization provided a testable rationale, since *ubK1* is repeatedly found in proximity to *porX* across *P. gingivalis* genomes, suggesting a functional pairing between the kinase and this orphan response regulator. PorX is a non-canonical response regulator that plays a central role in coordinating virulence-associated gene expression and secretion in *P. gingivalis* (17–19). Recent structural and functional studies have shown that PorX integrates inputs beyond canonical two component signaling, including Alkaline phosphatase domain (APS) associated oligonucleotide cleavage activity, and that it participates in regulatory circuits linked to Type-IX secretion system (T9SS) control (19,20). However, how PorX integrates these inputs and how they translate into specific downstream cellular outputs beyond gingipain based secretion readouts remain incompletely resolved. In this context, identifying additional upstream regulators of PorX is important for clarifying how PorX dependent signaling is wired within the broader *P. gingivalis* regulatory networks.

Here, we undertook a combined structural, biochemical, and genetic analysis to elucidate UbK1’s catalytic mechanism and place UbK1 within *P. gingivalis* regulatory networks. We present the crystal structure of UbK1 and use structure-guided mutagenesis to dissect its autophosphorylation mechanism, demonstrating that UbK1 can autophosphorylate in both *cis* and *trans*. Guided by conserved gene neighborhood analysis, we identify PorX as a previously unrecognized UbK1 substrate and map a single UbK1 dependent phosphosite in the PorX receiver domain. Finally, using a *ubK1* deletion strain in the W50 background and PorX phosphosite variant, we assess cellular outputs and show that UbK1 dependent PorX phosphorylation is separable from PorX canonical gingipain secretion output. Together, these findings establish a mechanistic framework for UbK1 function and expand UbK linked response regulator signaling in *P. gingivalis*.

## Methods

### Recombinant protein expression and purification in *Escherichia coli*

#### Construction of expression plasmids

Full length *porX* and *ubk1* genes were amplified from *Porphyromonas gingivalis* ATCC 33277 genomic DNA by Polymerase Chain Reaction (PCR). The PCR fragments were assembled into a modified pET28a(+) vector (Novagen) by Gibson assembly (21). In the modified expression vector, *porX* and *ubk1* genes were fused to N-terminal 10x-His tags followed by a Tobacco Etch Virus (TEV) protease recognition site or thrombin cleavage site, respectively. Site directed variants were generated by Gibson assembly using mutagenic primers. All primers and plasmids used in this study are listed in Supplementary Tables S1 and S2, respectively.

#### Bacterial cultivation and protein expression

Expression vectors were transformed into *E. coli* BL21(DE3) cells. Cultures were subsequently grown at 37°C in an autoinduction medium (22) containing kanamycin (50 μg/ml). After 8 hours, the temprature was shifted to 22°C for additional 16 hours. Cells were harvested by centrifugation at 6000 x g for 15 min at 4°C.

#### Recombinant protein purification

PorX and UbK1 (wild type and variants) were purified following a previously described method with slight modifications (23). Briefly, cell pellets were resuspended in buffer A (50 mM Tris pH 8, 300 mM NaCl and 10 mM imidazole) and incubated with DNase I (10 μg/ml) and protease inhibitor cocktail (Calbiochem) at 4°C. Cells were lysed using a French press pressure cell at 172 MPa. Cell debris was separated by ultracentrifugation at 270,000 × *g* for 1 hour at 4°C. Clarified supernatant was applied onto a gravity Ni-NTA column (Bio-Rad Econo-Column chromatography column, Thermo Scientific HisPur Ni-NTA resin) pre-equilibrated with buffer A. The bound protein was washed with buffers B (20 mM Tris pH 8, 300 mM NaCl and 20 mM imidazole) and C (20 mM Tris pH 8, 1 M NaCl and 40 mM imidazole) and eluted with buffer D (20 mM Tris pH 8, 200 mM NaCl and 500 mM imidazole). Affinity tags were removed by TEV for PorX or bovine thrombin (Prolytix) for Ubk1 digestion during an overnight dialysis against buffer E (20 mM Tris pH 8, 200 mM NaCl) at 4°C. Each protein was applied onto a size exclusion column (Superdex 200 16/60 GL, Cytivia) pre-equilibrated with buffer F (10 mM Tris pH 8, 200 mM NaCl). The purified proteins were then concentrated to ∼25 mg/ml and flash-frozen in liquid nitrogen.

### Crystallization and structure determination

Wild-type UbK1 was crystallized using the sitting drop vapour diffusion method at 25°C. A 0.3 μl solution containing 5 mg/ml UbK1, 1.25 mM BeSO_4_, 8.75 mM NaF and 630 μM MnCl_2_ was mixed with 0.3 μl of precipitant solution containing 0.1 M Bis Tris propane pH 6.33, 30% PEG 3350 and 0.2 M Na/K phosphate. Diffraction data were collected at the Canadian Light Source beamline CMCF-BM. 600 images were recorded at a wavelength of 1.18 Å with oscillation range of 0.2°, exposure time of 0.2 sec per image and a crystal-to-detector distance of 330.5 mm. Phases were obtained by molecular replacement in Phaser (24) with an Alphafold3 (25) model of UbK as an initial template. The final model was manually edited in Coot (26) and refined by Refmac5 (27). For R_free_ calculations, 5% of the data were excluded. All structural S6ures were prepared using ChimeraX (28). Data collection and refinement parameters are listed in Supplementary Table S3. Coordinates and structure factors have been deposited in the Protein Data Bank (accession code 9ZW5).

### Phosphotransfer assays

Thio-ATP labeling followed by alkylation and immunodetection was performed as described (6,29). For autophosphorylation reactions, 20 μg UbK1 was incubated with 500 μM ATP-γ-S in a kinase reaction buffer (25 mM Tris pH 7.5, 0.5 mM MgCl_2_, 0.5 MnCl_2_ and 25 μg/ml poly-L-lysine (PLL) at room temperature (RT) for 2 hours as described in prior UbK methods (6,13). For transphosphorylation, 20 μg PorX (either untreated, or from a 100 μM PorX pre-incubated with 20 mM Acetyl phosphate and 10 mM MgCl_2_ to induce REC domain dimerization or supplemented with 300 μM ZnCl_2_ to induce APS domain dimerization) was included in the reaction. Reactions were alkylated with 2 mM p-nitrobenzyl mesylate (PNBM) and incubated overnight at RT. The reactions were quenched with 5X loading dye containing 250 mM Tris pH 6.8, 10% SDS, 50% glycerol and 0.1% bromophenol blue, boiled at 95°C for 5 min and resolved by 20% SDS-PAGE. Proteins were subsequently electro-transferred onto a Polyvinylidene fluoride (PVDF) membrane pretreated with methanol and blocked for 1 hour at RT in a solution containing Tris-buffered saline with 0.1% Tween 20 (TBST) and 5% (w/v) skimmed milk. Thio-phosphorylated proteins were detected using a Rabbit monoclonal anti-thiophosphate ester antibody (Abcam) at a 1:5000 dilution in TBST supplemented with 5% skimmed milk, overnight 4°C. Membranes were washed three times with TBST before being probed for 60 min with a 1:10,000 dilution of a polyclonal goat anti-rabbit horseradish peroxidase-conjugated secondary antibody (Invitrogen) in TBST supplemented with 5% skimmed milk. Membranes were washed thrice with TBST and development was carried out using the enhanced chemiluminescence (ECL) western blot substrate kit according to the manufacturer’s instructions (Millipore).

### ATPase assays

ATP hydrolysis by UbK1 variants was quantified with the ADP-Glo Max Assay kit (Promega). Kinase reactions were first performed by incubating 10 μM UbK1 (WT or variants) with 500 μM ATP in a reaction buffer comprising of 25 mM Tris pH 7.5, 10 mM MgCl_2_, 1 mM MnCl_2_, and 2 mM dithiothreitol (DTT). Where indicated, 2.5 μM PorX was included in reactions containing WT UbK1. After incubation for 2 hours at RT, reactions were processed as per manufacturer instructions and luminescence signal was recorded by a BioTek Synergy plate reader. The reactions were carried out in quadruplicates.

### LC–MS/MS based phosphosite mapping

Autophosphorylation of UbK1 and subsequent transphosphorylation of PorX were carried out by incubating 50 μg of UbK1 with or without 50 μg PorX in a reaction buffer composed of 25 mM Tris pH 7.5, 0.5 mM MgCl_2_, 0.5 mM MnCl_2_, 0.2 mg/ml PLL and 1 mM ATP for 2 hours at RT in triplicates. A control reaction of PorX and ATP was set up in the absence of UbK1. The reactions were stopped by flash freezing in liquid nitrogen and stored at -80°C until further processing. For sample processing, reactions were diluted 2-fold in 10% SDS, 100 mM Triethylammonium bicarbonate (TEAB) pH 8.5, reduced with 10 mM Tris(2-carboxyethyl)phosphine (TCEP) at 95°C for 15 min, and alkylated with 40 mM iodoacetamide at RT in the dark for 15 min. An equivalent of 50 µg of protein was proteolytically digested with sequencing grade trypsin (Promega) at 37°C overnight using S-TRAP micro cartridges according to the vendor protocol. Tryptic peptides were lyophilized to dryness and phosphopeptides were enriched using zirconium immobilized metal affinity chromatography magnetic nanoparticles (Zr-IMAC HP, Resyn Biosciences). After enrichment, phosphopeptides were eluted in 1% ammonium hydroxide, neutralized by 5% Trifluroacetic acid and vacuum concentrated to dryness prior to rehydration in 0.1% formic acid. Samples were then loaded onto Evotip Pure tips (Evosep) according to the manufacturer protocol and analyzed using an Evosep One nanoLC and a Bruker timsTOF HT mass spectrometer operated in DDA-PASEF mode. Peptides were separated using the 30 sample per day (44 minute gradient, 500 nL/min) using an Evosep One nanoLC system and an EV1137 analytical column with a column heater set to 45°C. Mobile phases were a) 0.1 % formic acid in water, and b) 0.1% formic acid in acetonitrile. All solvents were of LC-MS purity. DDA-PASEF was conducted with MS survey scan (100 – 1800 m/z) and 10 PASEF ramps isolating the most abundant precursors for MS/MS fragmentation, with active exclusion of previously triggered precursors set to 0.4 min. PASEF intensity threshold and targets were set to 2500 and 20000 respectively. The ion mobility window was set between 0.7 and 1.4 1/K0. The CaptiveSpray source voltage was set to 1.6 kV. Collision energy was scaled from 20 – 59 eV between 0.6 and 1.6 1/K0. Raw data was searched, and precursor-based label free quantitation was performed using IonQuant (30) within Fragpipe version 23 (31). Database searching used reference sequences for PorX and UbK1 added into a FASTA file containing the canonical reference proteome for *E. coli* K12 strain (Uniprot, UP000000625, 4402 protein sequences). Methionine Oxidation (+15.9949 m/z), N-terminal acetylation (+42.0106), and S/T/Y phosphorylation (79.96633 m/z) were defined as variable modifications. Cleavage was set to strict trypsin with a maximum of 2 missed cleavages. Precursor and product m/z tolerances were set to 20 ppm. Differential phosphorylation analysis and visualization was conducted using Fragpipe Analyst (32) using both peptide and site-specific modification quantitation summaries as inputs. Features lacking quantitation in at least two-thirds of replicates within a given experimental condition were excluded. Site localization was further restricted by requiring that at least two of three replicates exhibit localization probabilities ≥0.9 in at least one condition. Significance was based on a Benjamini-Hochberg FDR adjusted p-value <0.05 and a Log2 fold-change of 1. Missing values were imputed using the perseus-type scheme in Fragpipe analyst. Data have been deposited to the ProteomeXchange Consortium via the PRIDE partner repository (33).

### Functional characterization in *P. gingivalis*

#### Bacterial strains, plasmids, and growth conditions

The wild-type strains used in this study was *P. gingivalis* W50. The wild-type bacteria and variants were propagated from -80°C freezer stocks and grown anaerobically at 37°C for 3-5 days on agar plates containing Trypticase Soy Broth (TSB) (Becton, Dickinson and Company, Franklin Lakes, NJ, USA) supplemented with 5 µg/ml hemin, 1 µg/ml menadione, 5% defibrinated sheep blood (BAPHK) (Hemostat Laboratories) and when required, 10 µg/ml Erythromycin or 1 µg/ml Tetracycline. The atmosphere of the anaerobic chamber contained a mixture of 5% hydrogen, 10% carbon dioxide and 85% nitrogen. The bacterial colonies were used to inoculate TSB broth supplemented with 5 μg/mL hemin and 1 μg/mL menadione, and cultures were grown anaerobically at 37°C without shaking.

#### Construction of deletion and single residues variants in P. gingivalis

The W50 deletion *ΔubK1* strain was generated as previously described. (19) Briefly, 1 kbp long regions both upstream and downstream of *ubK1*, were amplified from W50 genomic DNA and the erythromycin resistance gene (*emrF*) was amplified from plasmid pVA2198 by PCR. The three purified amplicons were assembled by Gibson assembly and transformed into *P. gingivalis* by electroporation. The transformed cells carrying the erythromycin resistance gene in the site of *ubK1* were selected by growth on BAPHK containing 10 μg/ml erythromycin. For PorX variant construction in *P. gingivalis* W50, our previously constructed *ΔporX* knockout strain (19) was used. The PorX point mutant Y79A was cloned into pT-COW under the *groES* constitutive promoter (PG0521) to generate pT-groES-PorX Y79A. From the same prior study, *ΔporX-*pT-COW (empty vector control) and *ΔporX*-pT-porX (wild type complement) were also used as controls. The pT-groES-PorX Y79A was introduced into *E. coli* S17-1 cells and transformed to W50 *ΔporX* by conjugation as previously described.(34) Briefly, BAPHK supplemented with tetracycline (1 μg/ml) was used to select pT-groES-PorX Y79A containing *P. gingivalis* cells and gentamicin (50 μg/ml) was used for the counter selection of the *E. coli* S17-1 donor cells. Transconjugants were obtained after 7 days of anaerobic incubation. They were then isolated and verified by PCR and Sanger sequencing. All primers and plasmids used in this study are listed in Supplementary Tables S1 and S2 respectively.

#### Gingipain enzymatic activity assay

Wild-type *P. gingivalis* W50, *ΔubK1,* W50 carrying empty pT-COW and *ΔporX* cells carrying empty pT-COW, pT-PorX and pT-PorX Y79A plasmids were cultivated to OD_600_ of 1.0. Cultures were harvested and screened for their gingipain activities. Arginine-gingipain (Rgp) and lysine-gingipain (Kgp) activities were measured as described previously (35) with minor modifications. For the Rgp assay, 10 μL of bacterial culture was mixed with 170 μL gingipain assay buffer (200 mM Tris HCl pH 7.6, 150 mM NaCl, 5 mM CaCl_2_, 0.02% NaN_3_ and 20 mM L-cysteine previously neutralized with 8 M NaOH in a 9:1 ratio). For the Kgp assay, 50 μL bacterial culture was mixed with 130 μL gingipain assay buffer. The mixtures were incubated at 37°C for 10 min followed by the addition of 1 mM substrate (Nα-benzoyl-L-arginine 4-nitroanilide hydrochloride for the Rgp assay and 2-acetamido-6-amino-*N*-(4-nitrophenyl) hexanamide for the Kgp assay). The formation of *p*-nitroaniline was measured at 405 nm at 1-min intervals for 30 min with constant shaking. All assays were performed using three biological replicates, each measured in four technical replicates.

#### Planktonic cell growth measurements

Wild-type W50 and *ΔubK1* W50 *P. gingivalis* strains were cultivated anaerobically at 37°C by inoculating cells from BAPHK plates into TSB media supplemented with 1 μg/mL menadione and varying concentrations of hemin (1 μg/mL, 5 μg/mL or 25 μg/ml). The cultures were incubated for 74 hours and OD600 was measured at defined intervals. Growth curves were performed with six independent biological replicates per condition.

## Results

### Structural characterization of UbK1

UbK1 was recombinantly overexpressed in *Escherichia coli* cells and purified to homogeneity as a monomeric protein (Fig. S1). Previous studies reported that UbK1’s phosphorylation activity shows a preference for manganese (Mn^2+^) over magnesium (Mg^2+^), in contrast to many of its homologs that are Mg^2+^ supported (13). Accordingly, the purified protein was screened for crystallization conditions designed to capture the different catalytic states of Ubk1, including the apo form, and complexes with nucleotides or their analogs in the presence of Mn^2+^ (ADP-Mn^2+^, AMP-PNP-Mn^2+^, BeF_3_^-^-Mn^2+^, ADP-AlF_3_-Mn^2+^ and ADP-VO_4_^3—^Mn^2+^). High resolution diffracting crystals were obtained exclusively under BeF_3_^-^-Mn^2+^ supplemented condition. UbK1 crystallized in the primitive orthorhombic space group (P2_1_2_1_2), with two monomers in the asymmetric unit (Fig. S2A). The two monomers are highly similar (Cα r.m.s.d = 1.04 Å), with localized confirmational heterogeneity in residue ranges of 7-16 and 53-63, where the maximum inter-monomer Cα displacements are 3.72 Å and 8.27 Å, respectively (Fig. S2B). Each Ubk1 monomer adopts the canonical UbK fold, consisting of a central β-sheet flanked by α-helices, consistent with previously characterized family members. The four conserved UbK signature motifs, including the Walker-A ATP binding motif (Gx4GKT), HxDxYR, SPT/S and EW motifs surround the central active site (Fig. 1A). Within the active site, K39 and T40 of the Walker-A motif coordinate a phosphate group and a Mn^2+^ cation. Although crystals were grown in the presence of BeF₃^-^, the additional electron density at the ATP-binding pocket is tetrahedral rather than trigonal planar and is best modeled as phosphate ion (Fig. S3). This assignment is further supported by the crystallization conditions, as the phosphate concentration in the drop (0.10 M from 0.20 M Na/K phosphate at 1:1 mixing) greatly exceeds the maximum possible BeF₃⁻ concentration (≤0.625 mM).

**Fig. 1.**
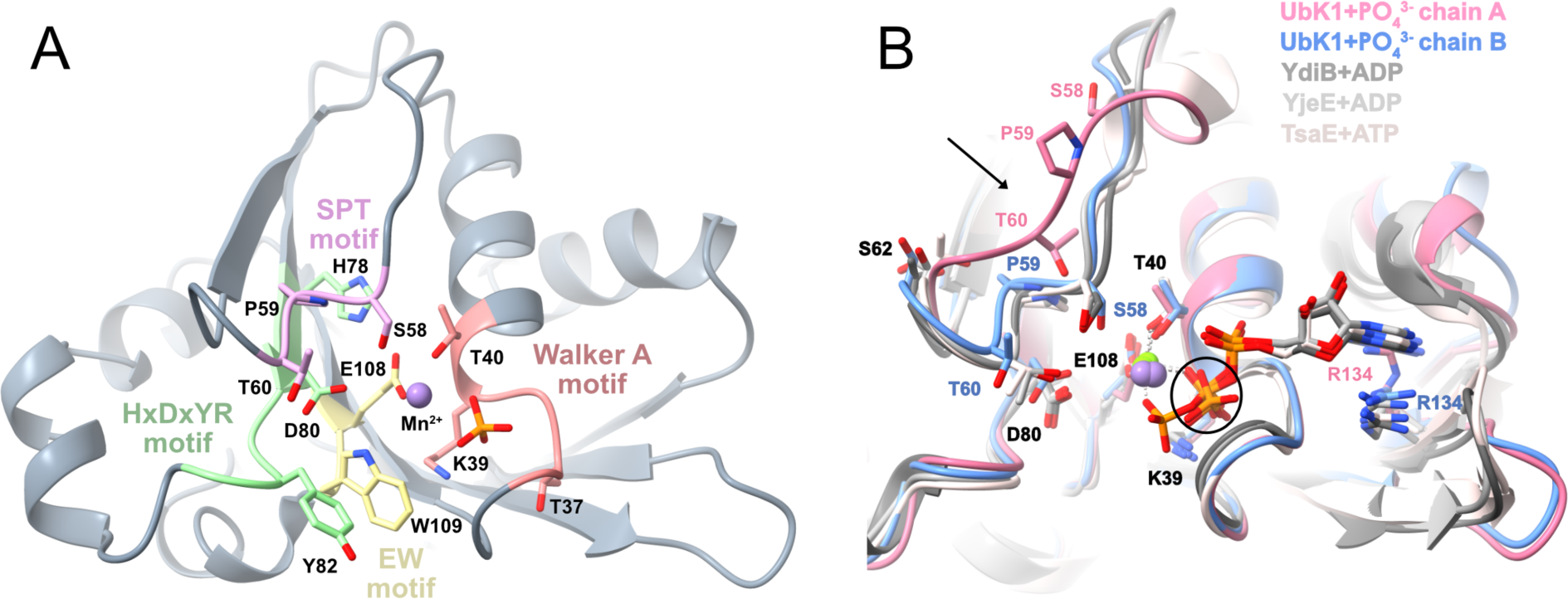
Structure of Ubk1 from *P. gingivalis.* A) UbK1 motif architecture highlighting conserved sequence elements surrounding the active site. B) Structural superposition of two UbK1 monomers in the asymmetric unit (pink and blue) with UbK homologs YdiB (PDB: 5NP9), YjeE (PDB: 1HTW), TsaE (PDB: 6N9A) shown in shades of grey. The overall fold is conserved, with localized structural differences most prominently in the SPT/S motif containing loop (black arrow). The black circle marks the β-phosphate observed in our crystal structure. Mn²⁺ and Mg²⁺ ions are shown as purple and green spheres, respectively.

The crystal structure of UbK1 shares high similarity with previously characterized homologs, including YdiB from *Bacillus subtilis*, YjeE from *Haemophilus influenzae* and TsaE from *Thermotoga maritima* (Ca r.m.s.d. = 1.78-1.82 Å, 1.8-1.85 Å, 2.19-1.85 Å, for chains A and B respectively). Structural superposition of UbK1 (bound to Mn^2^ and phosphate ion) with these nucleotide-bound homologs revealed conserved positioning of the Walker A motif relative to the β-phosphate and divalent metal cofactor. (6,14,36) In particular, the conserved lysine residue (K39_UbK1_) interacts with the β-phosphate and the conserved threonine residue (T40_UbK1_) coordinates the Mg^2+^/Mn^2+^ cations (Fig. 1B). The conserved glutamic acid residue of the EW motif (E108_Ubk1_) also directly coordinates the metal cation. Towards the ribose and adenine moieties of the nucleotide ligand in the three homologs structures, a conserved arginine is involved in van der Waals interactions with the adenine base (6). This residue is conserved in UbK1 (R134), suggesting a similar role. As the crystallization condition yielding diffracting crystals lacked nucleotide, the ribose and adenine-binding sites are unoccupied in our structure. In chain B, R134 adopts an alternative sidechain rotamer that occupies the pocket otherwise expected to accommodate the adenine base (Fig. 1B).

Within the UbK family, the HxDxYR motif in UbK family has been proposed to mirror the conserved catalytic HxD motif of the Hanks-type protein kinases, with the conserved aspartate (D80_UbK1_) positioned to coordinate the divalent metal cofactor and to facilitate transfer of the γ-phosphate to an acceptor residue on a protein substrate (6). The conserved histidine is thought to stabilize the local architecture surrounding the nucleotide, in part through interaction with E108 of the EW motif (6) (Fig. 1A). Consistent with structural and biochemical studies on YdiB and related UbKs, residues within and surrounding the SPT/S motif frequently undergo autophosphorylation and have been implicated in transphosphorylation of protein substrates (6,11). In UbK1, the principal conformational differences between the two monomers map to this region, which prior work indicates is intrinsically flexible (6). Superposition with homologous UbKs shows one UbK1 monomer adopting the canonical SPT conformation, whereas the other adopts a distinct arrangement (Fig. 1B). This flexibility is compatible with a role in autophosphorylation dynamics.

### Mechanistic analysis of UbK1 autophosphorylation

Members of the UbK family have been reported to hydrolyze ATP, undergo autophosphorylation and transphosphorylate protein substrates (6). Consistently, wild-type (WT) UbK1 from *P. gingivalis* exhibited ATP hydrolysis ability in an ATPase assay (Fig. 2A). Since Walker A containing proteins can in some cases bind and hydrolyze GTP (37), we also assessed UbK1 GTPase activity under identical conditions and could not detect GTP hydrolysis (Fig. S4). To monitor autophosphorylation, we employed a semi-synthetic immunoaffinity approach in which UbK1 was incubated with ATPγS to install thiophosphate on acceptor residues. These thiophosphorylated residues were subsequently alkylated with p-nitrobenzyl mesylate (PNBM) to generate stable thiophosphate ester epitopes, which were detected by immunoblotting by using a monoclonal anti-thiophosphate ester antibody.

**Fig. 2.**
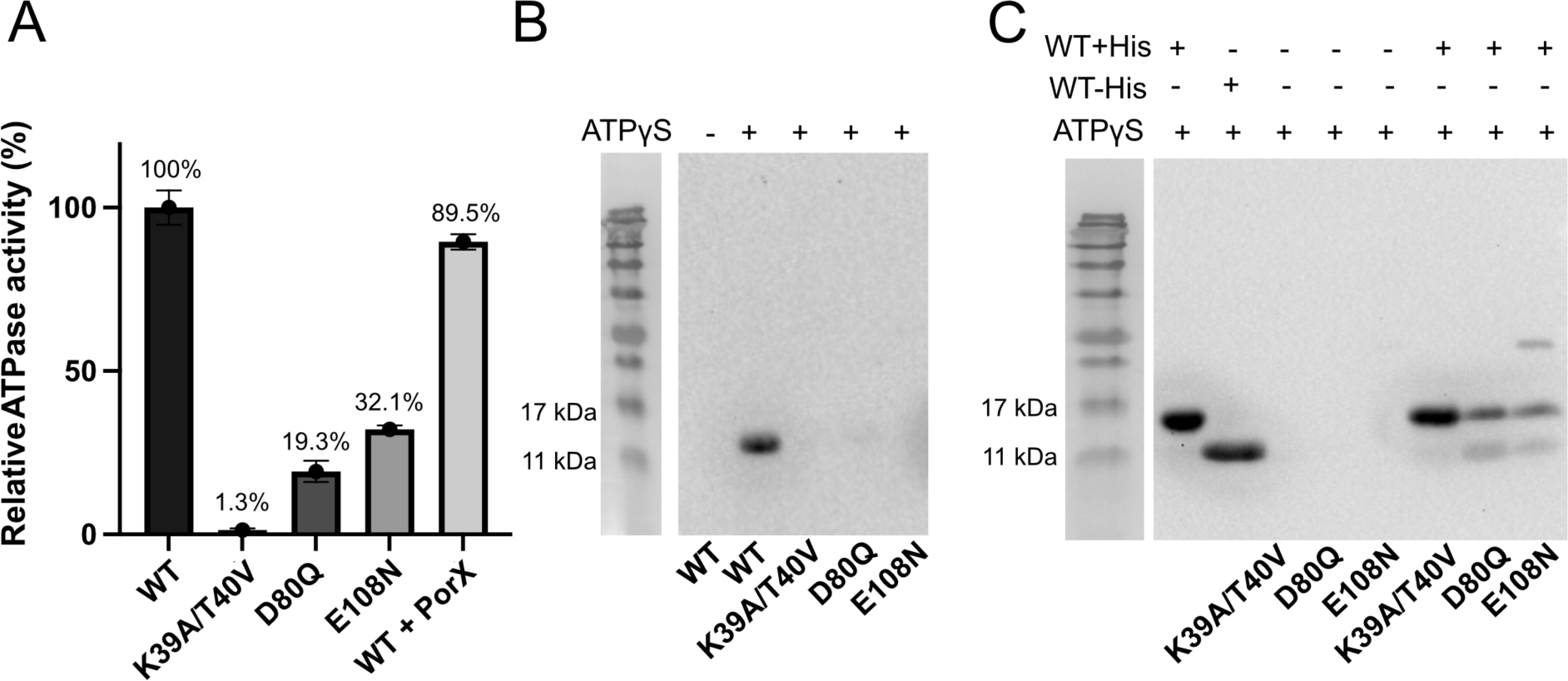
Catalytic activities and autophosphorylation mechanism of Ubk1 variants. A) ATPase activity of WT UbK1 with and without PorX and the indicated UbK1 variants. The reactions were carried out in quadruplicates and are reported relative to the corresponding WT control. B) ATPγS-dependent thiophosphorylation assay of WT UbK1 and the indicated variants. Mutations in the Walker A, HxDxYR, and EW motifs impair ATP hydrolysis and abolish or strongly reduce detectable autophosphorylation. C) Autophosphorylation mixing assay of His tagged WT UbK1 and tag-free catalytic variants (K39A/T40V, D80Q and E108N). Separation of His tagged and tag free species by SDS PAGE enables assessment of *cis* versus *trans* autophosphorylation and demonstrates UbK1 dependent phosphorylation in *trans* for D80Q and E108N.

To identify residues required for catalysis, we introduced structure-guided mutations in conserved motifs surrounding the ATP binding site. Disruption of the Walker A motif by K39A/T40V, which alters the β-phosphate contacting lysine and the metal coordinating threonine, reduced the ATPase activity to 1.3% of WT and eliminated detectable autophosphorylation (Fig. 2A-B). Mutations in the HxDxYR (D80Q) and EW motifs (E108N) reduced ATPase activity to 19% and 32% of the WT, respectively, and produced negligible autophosphorylation. (Fig. 2A-B) These observations are consistent with a role for D80 in facilitating γ-phosphate transfer to the acceptor residue and for E108 in coordinating the divalent metal cofactor. Together, these data establish essential roles for Walker A, HxDxYR, and EW motifs in ATP hydrolysis and kinase activities of UbK1.

Next, we preformed liquid chromatography-tandem mass spectrometry (LC-MS/MS) analysis to map the autophosphorylation sites in UbK1. In total, seven distinct phosphosites were identified in UbK1 (Fig. S5A, Extended Data). These include residues S53, T60 and S62 within and adjacent to the SPT motif; residues Y82 and Y77 within and adjacent to the HxDxYR motif; and residues S10 and Y67 at distal regions of the protein. In agreement with prior work in *P. gingivalis*, which identified S62, Y77 and Y82 as phosphosites, (13) our dataset confirms these sites and extends the UbK1 phosphosite map to a total seven residues, providing a more comprehensive view of UbK1’s autophosphorylation landscape.

Among the identified phosphosites, the highest-intensity phosphosignals localized to the SPT motif-containing loop, particularly at S62 and T60 (Extended Data), suggesting a potential regulatory role for this region. Notably, although multiple phosphorylatable residues were identified within and surrounding the SPT/S-containing loop (S53, T60, and S62), no UbK1-derived peptides carrying more than a single phosphorylation event within this region were detected in the LC–MS/MS analysis. All enriched phosphopeptides corresponding to the SPT/S loop region contained only one modified residue at a time. This observation indicates that autophosphorylation within the SPT/S loop does not result in stable accumulation of multiply phosphorylated loop species under the conditions tested, consistent with sequential or potentially mutually exclusive modification of individual residues. However, the absence of multiply phosphorylated SPT/S-loop peptides in the dataset does not exclude their transient formation, as multiply modified peptides are often underrepresented in shotgun phosphoproteomics due to reduced ionization efficiency, altered fragmentation behavior, and increased chromatographic heterogeneity (38,39). Consistent with this interpretation, the SPT/S loop displays pronounced conformational flexibility in the UbK1 crystal structure, which may limit simultaneous access of multiple residues to the catalytic site at any given time. Given the intrinsic flexibility of the SPT motif loop and its proximity to the D80 and γ-phosphate of the ATP moiety (docked into our UbK1 crystal structure), phosphosite mapping indicates that the majority of high-occupancy *cis*-autophosphorylation events localize to the SPT/S loop, while additional phosphorylation events at more distal sites likely arise through intermolecular phosphotransfer in *trans*.

To validate our hypothesis that UbK1 can in fact, undergo autophosphorylation in *trans,* in addition to *cis* as indicated by LCMS, we performed an assay whereby His-tagged WT UbK1 was mixed with catalytically deficient, tag-free UbK1 variants followed by ATPγS labeling and immunoblot analysis. The ∼2 KDa molecular weight difference between the His-tagged and tag-free proteins enables resolution by SDS-PAGE. For the Walker A double mutant (K39A/T40V), thiophosphorylation was detected exclusively on the His-tagged WT species (Fig. 2C), consistent with *cis*-only autophosphorylation, as previously reported for the *Streptococcus pneumoniae* UbK homolog (11). Notably, this Walker A double mutant represents the only residue combination previously used to investigate *cis* versus *trans* autophosphorylation within the UbK family. However, a strictly intramolecular autophosphorylation model is difficult to reconcile with our experimentally determined UbK1 structure and phosphosite mapping by mass spectrometry, which together are consistent with intermolecular autophosphorylation being feasible. We therefore extended this assay beyond the Walker A mutant to assess the effects of previously untested HxDxYR and EW motifs variants (D80Q or E108N) on autophosphorylation. In contrast to the Walker A mutant, both variants displayed clear trans-autophosphorylation signals (Fig. 2C). This differential behavior is most parsimoniously interpreted as reflecting differences in nucleotide-binding capacity among the mutants. In AAA ATPase family proteins, mutation of the conserved Walker A lysine, either alone or in combination with the adjacent threonine, has been shown to abolish nucleotide binding (40–43). By analogy, the K39A/T40V mutation in UbK1 is expected to severely impair or eliminate nucleotide binding, which could prevent this variant from engaging in ATP-dependent phosphorylation reactions. While alternative explanations are possible, differences in nucleotide-binding capacity provide a straightforward interpretation consistent with prior work.

More generally, nucleotide binding is thought to represent an early step in kinase catalysis and may be required to promote a conformation competent for productive interaction with a phosphorylation substrate. From this perspective, loss of nucleotide binding in the Walker A mutant could plausibly preclude its phosphorylation by WT UbK1 in trans, independent of its intrinsic catalytic potential. In contrast, the D80Q and E108N variants are predicted to retain nucleotide-binding capability despite impaired ATP hydrolysis, (40) potentially allowing them to adopt a nucleotide-bound state that can be recognized and phosphorylated by WT UbK1. Taken together, these results support the conclusion that UbK1 is capable of autophosphorylation in *trans*. While *cis*-autophosphorylation also occurs, our findings argue against a strictly intramolecular autokinase mechanism for UbK1. Establishing whether UbK1 can act intermolecularly is important for understanding how this kinase may engage and phosphorylate external protein substrates.

### Transphosphorylation of response regulator PorX by UbK1 *in vitro*

Previous studies in *P. gingivalis* identified the response regulator RprY as a substrate of UbK1 (16). Given the relaxed substrate specificity reported for UbK homologs and the prevalence of orphan response regulators in *P. gingivalis* signaling networks, we reasoned that UbK1 may engage additional protein substrates. Since bacterial signaling components are frequently organized into conserved local gene neighborhoods, we used operon conservation as a guide to nominate additional candidate substrates. To this end, we examined the genetic neighborhood of *ubK1* across 81 *P. gingivalis* strains with available genome sequences. This analysis revealed that *ubK1* is predicted to reside in a conserved three gene operon that includes *porX*, encoding another orphan response regulator, and *imm17,* predicted to encode an immunity factor protein. Based on this genomic organization, we hypothesized that PorX might represent an additional UbK1 substrate. In *in vitro* phosphotransfer assays containing UbK1 and PorX, UbK1 catalyzed robust transphosphorylation of PorX (Fig. 3A). In contrast, PorX alone displayed only weak background ATPγS dependent labeling (Fig. 3B). Catalytically impaired UbK1 variants (K39A/T40V, D80Q, and E108N) failed to increase PorX labeling above the PorX-alone baseline (Fig. 3A-B), indicating that PorX transphosphorylation depends on UbK1 catalytic activity rather than on a passive UbK1-PorX association.

**Fig. 3.**
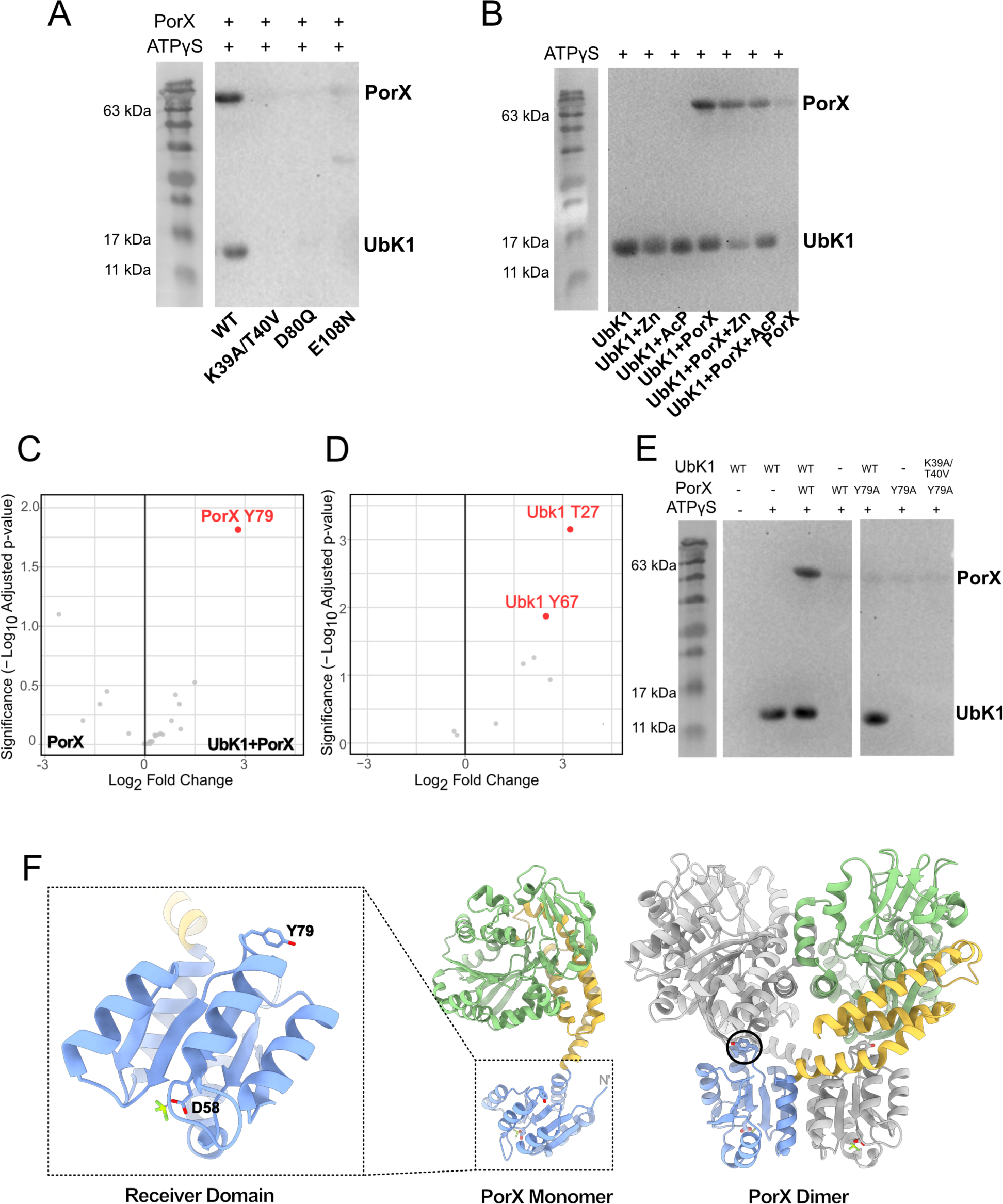
UbK1 transphosphorylates PorX at Y79 independently of PorX oligomeric state. A) ATPγS dependent phosphotransfer assay showing UbK1 autophosphorylation and subsequent PorX transphosphorylation. B) Phosphotransfer assays using monomeric PorX and PorX dimers induced by acetyl phosphate or Zn²⁺ binding show that UbK1 transphosphorylates PorX under all oligomeric conditions tested. C) Volcano plot from LC MS/MS differential phosphorylation analysis comparing UbK1 incubated with PorX vs PorX alone, demonstrating significant enrichment of phosphorylation at PorX Y79 in the presence of UbK1. D) Volcano plot from LC MS/MS differential phosphorylation analysis comparing UbK1 incubated with PorX vs UbK1 alone, exhibiting phospho-enrichment at UbK1 Y67 and addition of a new site; UbK1 T27 in the presence of PorX. E) Phosphotransfer assay validating Y79 as the primary UbK1 dependent PorX phosphosite. The Y79A substitution reduces UbK1 dependent PorX labeling to the baseline level observed for PorX alone. F) Mapping of Y79 onto PorX’s structure (PDB: 7PV7) shows that Y79 resides in PorX’s receiver (REC) domain. Black circle indicates Y79 position on PorX dimer.

We next examined whether the oligomeric state of PorX affects UbK1-dependent transphosphorylation. Our previous structural and functional characterization of PorX demonstrated that monomeric PorX contains a receiver (REC) domain and an alkaline phosphatase (APS) domain connected by a three-helix bundle (19,20). Upon activation induced dimerization, two PorX monomers form an intertwined X-shaped assembly with dimer interfaces at the REC and the APS domains. Dimerization of PorX can be induced by phosphorylation of the REC domain with acetyl phosphate (AcP) or by zinc binding at the APS phosphatase site (19,20). We compared UbK1-dependent transphosphorylation of PorX in its monomeric state and after dimerization with AcP or Zn^2+^. UbK1 transphosphorylated PorX under all three conditions with no evident preference for oligomeric state, which suggests that the transphosphorylated residue or residues are not protected by the dimer interfaces and are accessible in both oligomeric states. Control reactions confirmed that AcP and Zn^2+^ did not affect UbK1’s autophosphorylation activity (Fig. 3B).

To further characterize the UbK1–PorX phosphorylation relationship, we performed LC–MS/MS differential phosphorylation analysis to assess pattern changes upon incubation of the two proteins. Consistent with the phosphotransfer assays, PorX incubated with UbK1displayed significant enrichment of phosphorylation at a single residue, Y79 (Fig. 3C, Extended Data). In contrast, PorX incubated with ATP alone exhibited only low-intensity phosphorylation at multiple sites, consistent with the weak background signal observed in the PorX-only lane of the phosphotransfer assay immunoblot (Fig. 3B, Extended data). Importantly, these additional sites were not significantly enriched upon incubation with UbK1, indicating that they likely reflect background or UbK1-independent phosphorylation events (20).

In parallel, UbK1 exhibited largely stable autophosphorylation patterns in the presence of PorX. However, differential phosphorylation analysis revealed increased phosphorylation at two UbK1 residues, Y67 and T27, in the UbK1+PorX condition (Fig. 3D, Extended Data 1). Phosphorylation at Y67 was detected previously in UbK1 alone, whereas T27 represents a newly observed phosphosite that was not detected in the UbK1-only condition. Notably, both residues are located distal to the catalytic center and the SPT/S loop, making intramolecular *cis*-autophosphorylation unlikely (Fig. S5A). These observations are therefore consistent with phosphorylation events arising through intermolecular phosphotransfer in *trans*. It is likely that phosphorylation of the SPT/S motif might cause conformational changes making Y67 more accessible for intermolecular phosphotransfer to occur, thereby causing an enrichment of this phosphosite. Despite these site-specific changes, PorX addition did not detectably alter UbK1 ATP hydrolysis under the assay conditions used (Fig. 2A), indicating that PorX does not globally stimulate or inhibit UbK1 ATPase activity.

To validate the site assignment of Y79 on PorX, we generated and tested a PorX Y79A mutant in the phosphotransfer assay. This single Y79A substitution markedly reduced UbK1-dependent PorX transphosphorylation to a level comparable to PorX alone, supporting Y79 as the primary UbK1-dependent phosphorylation site (Fig. 3E). Mapping Y79 on the PorX determined structure revealed that this residue resides within the receiver (REC) domain and is spatially distinct from the conserved D58, which serves as the canonical phosphorylation site in response regulators. Together, these data identify Y79 as a previously unrecognized regulatory site on PorX (Fig. 3F).

### *In vivo* functional characterization of UbK1 suggests PorX regulatory roles beyond the Type-IX secretion system

Previous work reported *ubK1* as an essential gene in *P. gingivalis* strain ATCC 33277, and functional analyses were therefore limited to overexpression-based approaches in that background (13). Here we successfully constructed a *ubk1* deletion variant in the W50 strain. Notably, W50 strain differs from ATCC 33277 strain in several virulence-relevant traits, including the ability to synthesize a protective capsule, reduced fimbriae expression, altered auto-aggregation, and enhanced virulence phenotypes in animal models (44–46). These strain-specific differences, together with additional genetic or regulatory factors, may contribute to the apparent essentiality in *ubk1* in one strain background but not the other.

Earlier work showed that PorX protects *P. gingivalis* 33277 strain cells from hemin overload by upregulating specific gene expression programs (47). Given our identification of PorX as a UbK1 substrate, we next examined whether UbK1 contributes to a similar protective response. Growth of W50 WT and W50 *Δubk1* strains was compared across several hemin concentrations. Loss of *ubk1* consistently slowed bacterial growth with the deletion strain failing to reach the same maximal cell density as the WT strain (Fig. 4A). However, growth defects were comparable across all hemin conditions tested, indicating that UbK1 does not play a major role in protection against hemin overload.

**Fig. 4.**
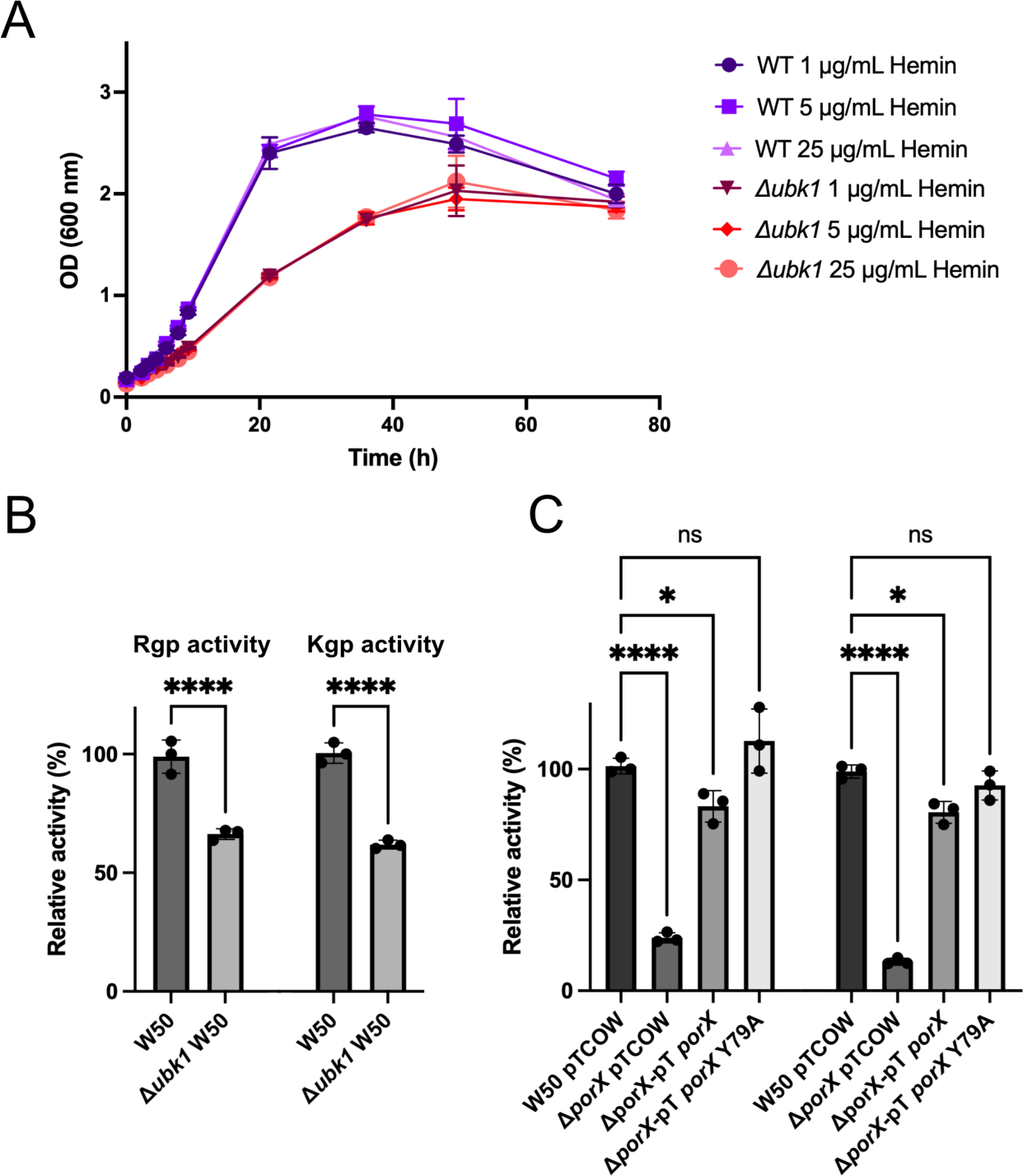
*In vivo* phenotypes of *ubK1* deletion and functional separation of UbK1-dependent PorX phosphorylation from gingipain secretion. A) Growth curves of WT and *Δubk1* across hemin conditions show that *ubk1* deletion reduces growth rate and maximal cell density, with comparable defects across the conditions tested. Growth curves were performed with six independent biological replicates per condition. (B) Gingipain activity assays (Rgp and Kgp) comparing WT and *Δubk1* strains show a moderate reduction in extracellular gingipain activity upon *ubK1* deletion. C) Gingipain activity assays in *ΔporX* complemented strains including empty plasmid complementation, WT complementation (*ΔporX* pT-PorX) and the transphosphorylation site variant (*ΔporX* pT-PorX Y79A). PorX Y79A shows gingipain activity comparable to WT and the complemented strain, indicating that UbK1 dependent phosphorylation of PorX at Y79 is dispensable for T9SS dependent gingipain secretion under the conditions tested. Gingipain activity assays were performed using three biological replicates, each measured in four technical replicates. Statistical significance was assessed by one way ANOVA followed by Šídák’s multiple comparisons test with adjusted p-values <0.05 considered significant.

In addition to its role in regulating gene expression in response to hemin, PorX is a key regulator of T9SS, a specialized secretion pathway required for the maturation and extracellular export of major virulence factors in *P. gingivalis* (17,18,47). Gingipains which are arginine and lysine specific cysteine proteases, are the best-characterized substrates of the T9SS, and their extracellular activity is widely used as a functional proxy for T9SS assembly and secretion efficiency (18,19,48). Deletion of *porX* has been shown to cause a severe defect in gingipain secretion due to impaired T9SS function (19). Since PorX is phosphorylated by UbK1, we therefore asked whether loss of *ubk1* similarly affects gingipain secretion, which would suggest that UbK1 contributes to PorX dependent regulation of the T9SS. Using gingipain activity assays, we observed a moderate decrease in Rgp/Kgp secretion, although this effect was substantially less pronounced than that caused by *porX* deletion (Fig. 4B). While these results suggest a possible contribution of UbK1 to gingipain secretion, they do not provide strong evidence that UbK1 regulates gingipain export directly through the PorX regulatory cascade. To directly assess whether the UbK1-meidated phosphorylation of PorX is functionally linked to gingipain secretion, we measured gingipain activity in the PorX Y79A variant. This variant exhibited gingipain activity comparable to that of the wild-type protein (Fig. 4C), indicating that UbK1-mediated phosphorylation of PorX at Y79 is unlikely to play a major role in T9SS-dependent gingipain secretion. Together, these findings argue that UbK1-mediated phosphorylation of PorX regulates functions distinct from its role in T9SS-dependent gingipain secretion.

## Discussion

The UbK family represents a broadly conserved yet still incompletely understood class of bacterial protein kinases with dual specificity toward serine/threonine and tyrosine residues. Although UbK homologs are present across diverse bacterial phyla, only five have been functionally characterized as kinases, and even fewer (=3) have been examined in structural detail (6,11–14,36). Existing studies link UbK family members to a range of cellular processes, including translation-associated pathways (via t^6^A-related machinery), growth control, stress tolerance, and virulence-associated behaviors (6,10,11,14,49). However, how UbK kinases integrate into broader signaling networks, how conserved motifs contribute to catalysis, and how autophosphorylation is coordinated with substrate phosphorylation remain open questions.

In this study, we provide a combined structural, biochemical, and genetic analysis of UbK1 from *Porphyromonas gingivalis*, advancing mechanistic understanding of UbK family kinases and clarifying UbK1’s relationship to orphan response regulator signaling in this organism. The crystal structure of UbK1 adopts the canonical UbK fold closely resembling previously characterized homologs such as YdiB, YjeE and TsaE (Fig. 1A-B) (6,11,14,36) and positions the conserved Walker A, HxDxYR, SPT/S and EW motifs around the ATP-binding pocket (Fig. 1A). Structure-guided mutagenesis confirms the Walker A, HxDxYR and EW motifs are essential for ATP hydrolysis and kinase activity, consistent with prior biochemical analyses reported for the YdiB and UbK *(S. pneumoniae)* homologs (6,11) (Fig. 2A-B). Notably, UbK1 exhibits pronounced conformational heterogeneity in the SPT/S motif region, directly visualizing the flexibility previously inferred for this loop (6) and suggesting that the SPT/S region can sample multiple states during UbK-mediated phosphorylation (Fig. 1B).

### Autophosphorylation architecture: cis hotspot in the SPT/S loop and distal phosphorylation in trans

Across characterized UbK homologs, ATPase activity is often low, yet autophosphorylation is consistently observed and accompanies phosphotransfer to protein substrates (6,50). This pattern suggests that autophosphorylation is a recurring component of UbK enzymatic behavior rather than a rare or purely regulatory event, and it is often discussed in a different context from the activation focused autophosphorylation that is commonly highlighted for Hanks type kinases (51,52). Consistent with this framework, UbK1 hydrolyzes ATP and undergoes robust ATPγS dependent thiophosphorylation, and disruption of conserved catalytic motifs abolishes both ATPase activity and autophosphorylation, supporting a functional coupling between nucleotide processing and phosphorylation activity (Fig. 2). LC–MS/MS phosphosite mapping further expands the known UbK1 phosphorylation landscape and provides insight into how phosphorylation is distributed across the protein. In UbK1 incubated with ATP alone, seven phosphosites were detected with high confidence, spanning the SPT/S region and additional distal positions (Fig. S5A). The most prominent phosphosignals localized to the SPT/S loop, particularly at T60 and S62, supporting the view that this conformationally flexible loop is a primary autophosphorylation hotspot (Extended data).

Importantly, although multiple phosphorylatable residues reside within and adjacent to the SPT/S loop, we did not detect UbK1 derived phosphopeptides carrying more than one phosphorylation event within this region under the conditions tested. This suggests that UbK1 does not stably accumulate multiply phosphorylated SPT/S loop species, which is consistent with sequential or mutually exclusive modification of individual residues. This conclusion should be interpreted cautiously because multiply modified peptides are often underrepresented in shotgun phosphoproteomics due to reduced ionization efficiency, altered fragmentation behavior, and increased chromatographic heterogeneity (38,39). In addition, the pronounced conformational flexibility of the SPT/S loop observed in the UbK1 crystal structure may limit simultaneous access of multiple residues to the catalytic site at any given time, thereby favoring transient or sequential modification rather than stable multi-site loop phosphorylation.

Together, these observations support a working model in which *cis* autophosphorylation predominates within the SPT/S loop, whereas phosphorylation events at more distal sites are more consistent with intermolecular phosphotransfer in trans. This division is conceptually useful because it helps reconcile how UbK kinases can preserve a conserved catalytic core while producing diverse phosphorylation signatures across species. UbK family members nevertheless display substantial diversity in the number and identity of autophosphorylation sites (Fig. S5B). For example, YdiB from *Bacillus subtilis* accumulates four S/T sites centered on and flanking the SPT/S motif (T55, S60, T62, T64) (6), whereas the *Streptococcus pneumoniae* UbK homolog was reported to autophosphorylate predominantly at a single tyrosine within the SPT/S region (Y58) (11). In *Streptococcus pyogenes*, the SP-TyK homolog carries two autophosphorylated tyrosine residues spanning the SPT/S region and the HxDxYR-associated segment (Y59 and Y77) (12). In *P. gingivalis*, UbK1 was previously reported to autophosphorylate at S62, Y77, and Y82 (13), and our phosphosite mapping corroborates these sites while expanding the UbK1 autophosphorylation landscape. Despite this diversity, all UbK kinases characterized to date share at least one autophosphorylation site within or adjacent to the SPT/S loop, supporting this region as a recurrent hotspot for UbK autophosphorylation.

Finally, differential phosphorylation analysis in the presence of PorX further supports the idea that distal phosphorylation can arise through intermolecular transfer. In this condition, phosphorylation at Y67 increased and an additional UbK1 phosphosite at T27 became detectable, whereas UbK1 incubated alone did not show phosphorylation at T27. Both residues are located distal to the active site and the SPT/S loop, which is consistent with phosphorylation arising through intermolecular phosphotransfer rather than intramolecular loop centered *cis* modification.

### UbK1 undergoes both cis- and trans-autophosphorylation: nucleotide binding as a gate for intermolecular phosphorylation

Mechanistically, our mixed-protein assays demonstrate that UbK1 can autophosphorylate in both *cis* and *trans* (Fig. 2C). The Walker A double mutant (K39A/T40V) shows thiophosphorylation only on the His-tagged wild-type species, consistent with *cis*-only autophosphorylation in that assay configuration, as previously reported using a comparable Walker A mutant in the *S. pneumoniae* UbK homolog (11). However, extending the assay beyond Walker A revealed a different outcome with the catalytically impaired HxDxYR (D80Q) and EW (E108N) variants displaying a clear phosphorylation in *trans* when mixed with wild-type UbK1 (Fig. 2C). To our knowledge, this provides direct evidence that UbK family kinases are not restricted to a strictly intramolecular autokinase mechanism. It also underscores that conclusions about *cis* only autophosphorylation can depend on the specific mutant framework used to probe intermolecular phosphotransfer, since Walker A double mutants can severely compromise nucleotide binding and therefore may be poorly suited for testing *trans* autophosphorylation.

A parsimonious interpretation is that nucleotide-binding capacity acts as a prerequisite for productive intermolecular phosphorylation, even when catalytic turnover is compromised. The Walker A double mutant is expected to severely impair or eliminate nucleotide binding, which could prevent it from adopting the conformation required for productive substrate engagement and thus preclude its phosphorylation by wild-type UbK1 in *trans*. In contrast, D80Q and E108N are more likely to retain nucleotide binding while being defective in catalytic chemistry, enabling them to engage wild-type UbK1 and become phosphorylated in *trans*. While alternative explanations are possible, differences in nucleotide-binding capacity provide a straightforward interpretation consistent with prior work.

Together, these data support a model in which *cis* autophosphorylation can occur, but UbK1 is also capable of intermolecular phosphotransfer. Establishing that UbK1 can phosphorylate in *trans* is important for understanding how this kinase may engage and phosphorylate external protein substrates.

### PorX is a UbK1 substrate: specificity for PorX Y79 and independence from PorX oligomeric state

Beyond autophosphorylation, UbK kinases function through transphosphorylation of protein substrates. UbK1 was previously shown to phosphorylate the orphan response regulator RprY, linking UbK signaling to community development and virulence-associated behaviors in *P. gingivalis* (13). Here, guided by conserved gene neighborhood analysis, we identify PorX as a second UbK1 substrate. UbK1 robustly transphosphorylates PorX *in vitro*, (Fig. 3A-B) and this phosphorylation depends on UbK1 catalytic function rather than reflecting passive association.

Differential phosphoproteomics indicates that UbK1-dependent phosphorylation of PorX is highly specific. A single PorX residue is significantly enriched upon incubation with UbK1, Y79 in the receiver domain (Fig. 3E, Extended data). By comparison, PorX incubated with ATP alone displays only low intensity phosphorylation at multiple sites, consistent with the weak background signal observed in PorX only phosphotransfer conditions. Importantly, these additional sites are not enriched in the presence of UbK1 (Extended data), supporting the interpretation that they reflect background or UbK1 independent events. Functional testing supports the proteomics assignment, as the PorX Y79A substitution reduces UbK1-dependent PorX labeling to a level comparable to PorX alone, consistent with Y79 being the primary UbK1-dependent site under these conditions.

UbK1 phosphorylates PorX across all oligomeric conditions tested. UbK1 transphosphorylates monomeric PorX and PorX dimers induced by acetyl phosphate or zinc binding. This indicates that the UbK1 target site remains accessible in both monomeric and dimeric PorX states and is not occluded by the characterized dimer interfaces (Fig. 3F).

Comparison with RprY further supports substrate specificity. UbK1 targets different tyrosine residues in PorX and RprY despite their shared receiver domain fold, which supports a model in which UbK1 recognition depends on substrate specific features rather than phosphorylation of a single conserved receiver domain position. PorX Y79 residue is not broadly conserved across T9SS-containing bacteria, consistent with the idea that UbK substrate interfaces and downstream consequences may be species specific. This point is especially relevant because PorX homologs contribute to T9SS control across the Bacteroidota, yet the dominant biological outputs coupled to T9SS activity differ substantially between organisms. In *P. gingivalis*, PorX is tightly linked to secretion of major virulence factors, whereas in other species PorX related regulatory modules have been associated with more modest effects on secretion of metabolic enzymes and no clear role in gliding motility in *Flavobacterium johnsoniae* (19). Together with our finding that PorX Y79 phosphorylation is dispensable for gingipain secretion, these observations are consistent with our previous suggested model that PorX functions as a regulatory hub in *P. gingivalis*, integrating inputs that influence pathways beyond T9SS dependent cargo export (19,20).

### Substrate engagement modestly alters UbK1 phosphorylation: Y67 and PorX-induced T27 point to trans-phosphorylation events

An additional nuance emerging from differential phosphorylation analysis is that incubation with PorX does not broadly remodel UbK1’s autophosphorylation landscape, but it does increase phosphorylation at two UbK1 residues: Y67 and T27, with T27 appearing as a PorX-associated site not detected in UbK1 alone. Both residues map distal to the active site and SPT/S loop, making intramolecular *cis*-autophosphorylation unlikely. A conservative interpretation is that substrate engagement can promote additional intermolecular phosphorylation events in UbK1, either through transient protein-protein interaction or through altered conformational sampling that enables *trans*-autophosphorylation at distal positions. Whether these modifications contribute to signaling output or reflect “non-productive” enzyme cycling remains unresolved. Importantly, because these changes are limited and site-specific, they support a model in which UbK1–PorX interaction subtly influences UbK1 phosphorylation state without globally altering UbK1 activity, consistent with the observation that UbK1 ATPase output does not change detectably upon addition of PorX under the conditions tested (Fig. 2A).

### In vivo phenotypes support PorX roles beyond gingipain secretion and indicate UbK1–PorX signaling is separable from canonical T9SS outputs

To probe UbK1 function in the bacterial context, we generated a ubk1 deletion strain in the *P. gingivalis* W50 background. This contrasts with prior work in strain ATCC 33277, in which *ubk1* could not be deleted (13). The W50 and ATCC 33277 backgrounds differ in several virulence relevant traits, including capsule production, fimbrial phenotypes, aggregation behavior, and infection associated phenotypes, (44–46) and these differences could plausibly influence the observed essentiality constraints. Additional genetic or regulatory factors are also likely to contribute.

Loss of *ubk1* produced a reproducible growth defect, with reduced growth rate and a lower maximal cell density (Fig. 4A). This phenotype is consistent with a conserved contribution of UbK family kinases to bacterial fitness. Growth or fitness defects following UbK perturbation have been reported for homologs in *Streptococcus pneumoniae* and *Streptococcus mutans*, among other species (6,10,11), indicating that this phenotype is not unique to *P. gingivalis*. Importantly, in our assays the magnitude of the growth defect was similar across the hemin concentrations tested, indicating that UbK1 does not play a major role in protection against hemin overload under these conditions. This distinguishes UbK1 from PorX, which has been implicated in hemin responsive gene regulation (53), and suggests that UbK1 contributes to cellular fitness through pathways not captured by hemin overload phenotyping.

Since PorX is required for efficient T9SS-dependent export of major virulence factors in *P.gingivalis*, we next tested whether UbK1 impacts gingipain secretion. Deletion of *ubk1* caused a moderate decrease in extracellular gingipain activity (Fig. 4B), but the effect was substantially weaker than the severe gingipain secretion defect associated with *porX* deletion (19). Critically, the PorX Y79A variant, despite being strongly defective for UbK1-dependent phosphorylation *in vitro*, exhibited gingipain activity comparable to wild-type PorX (Fig. 4C). This indicates that UbK1-dependent phosphorylation at Y79 is not required for PorX’s support of gingipain secretion.

Together, these *in vivo* results support the conclusion that UbK1-PorX signaling is functionally separable from PorX’s canonical role in T9SS dependent gingipain secretion. This, in turn, is consistent with prior evidence that PorX participates in additional signaling processes beyond gingipain export, including APS domain mediated cleavage of cyclic oligoadenylates, an activity that is dispensable for gingipain secretion. Collectively, these observations support a model in which PorX functions as a multifunctional signaling hub in *P. gingivalis*, integrating inputs that influence regulatory outputs beyond T9SS-dependent cargo export.

### Interpreting the modest gingipain phenotype: indirect effects and network-level positioning

The modest gingipain defect in Δ*ubK1* is more consistent with indirect effects rather than direct control of the T9SS through PorX Y79 phosphorylation. One plausible route is via RprY-dependent regulation, since *rpry* deletion has been reported to alter expression of multiple virulence-associated factors, including components linked to gingipain output (15). If UbK1 acts upstream of more than one orphan response regulator, then loss of UbK1 could shift regulatory states in a way that modestly reduces gingipain activity without directly perturbing PorX functions required for T9SS dependent cargo export.

To place UbK1 relative to the T9SS machinery, we reanalyzed a published label free proteomics dataset of *P. gingivalis* T9SS gene deletions (54). In that dataset, PorX abundance changed markedly across multiple T9SS knockouts, including *ΔporU, ΔporV, ΔporZ, ΔporT*, and *ΔporE*, consistent with tight coupling between PorX and T9SS component status. In contrast, UbK1 did not show a consistent correlation with T9SS gene deletions in the same comparisons (Fig. S6). Although most of these mutants were generated in strain ATCC 33277, the core T9SS machinery is conserved across *P. gingivalis* strains, and key protein dependencies within the apparatus are broadly shared (17–19,47,54–56). Together with our observation that *ΔubK1* causes only a moderate reduction in gingipain secretion and that PorX Y79A does not impair gingipain output, this pattern supports a model in which UbK1 contributes to secretion associated phenotypes indirectly, rather than functioning as a dedicated component of the PorX dependent T9SS regulatory axis.

More broadly, the fact that the only two UbK1 substrates identified to date, RprY and PorX, are orphan response regulators with established links to virulence associated pathways suggests that UbK1 may intersect with multiple virulence relevant circuits, alongside its more general contribution to cellular fitness reported for other UbK homologs (6,10,11).

Although our data establish UbK1-dependent phosphorylation of PorX, the downstream functional consequences of this modification remain unresolved, in part because we currently lack an assay that can isolate the specific effects of Y79 phosphorylation from broader network-level outputs. We therefore tested two biologically motivated readouts linked to established PorX functions, protection against hemin overload (47) and PorX-dependent gingipain secretion through the T9SS (19), but neither assay provided a definitive correlate of PorX Y79 phosphorylation. In particular, *ubk1* deletion produced a modest gingipain phenotype, whereas PorX Y79A did not measurably alter gingipain secretion, indicating that this modification is not required for the canonical PorX output measured by this assay.

Resolving the consequences of UbK1-PorX phosphorylation will likely require systems-level approaches that can capture pathway-level rewiring rather than single-output phenotypes. Comparative whole-cell proteomics and transcriptomic profiling of *ubK1* and *porX* deletion strains can define the broader regulatory networks in which UbK1 and PorX operate and provide an unbiased route to identify candidate pathways and regulatory partners. These datasets would also enable systematic expansion of the UbK1 and PorX protein interactomes and help prioritize modules relevant to host interaction and virulence. Such systems-level analyses and subsequent functional validation of nominated pathways lie beyond the scope of the present study and represent an important direction for future work.

### Conclusions

In summary, we establish a structural and mechanistic framework for UbK1 and define key conserved motifs required for ATP hydrolysis and phosphorylation activity. We demonstrate that UbK1 undergoes autophosphorylation in both *cis* and *trans* supporting a model in which nucleotide binding enables productive intermolecular phosphotransfer. We further identify PorX as a previously unrecognized UbK1 substrate and map a highly specific UbK1-dependent phosphosite on PorX (Y79). Although PorX Y79 phosphorylation is dispensable for gingipain secretion, these data expand the UbK substrate repertoire and support a model in which UbK1 interfaces with orphan response regulator signaling pathways that extend beyond canonical T9SS-associated outputs in *P. gingivalis*.

## Data availability

Atomic model and structure factors files have been deposited in the Protein Data Bank (PDB); accessions code: 9ZW5.

The mass spectrometry proteomics data have been deposited to the ProteomeXchange Consortium via the PRIDE partner repository with the dataset identifier PXD072953.

## Acknowledgments

We would like to thank Dr. Vincent Richard and Dr. Timon Geib at the Clinical Proteomics Center at the Lady Davis Institute, McGill University for support with LCMS/MS experiments. We would also like to extend our gratitude to the Canadian Light Source (CLS) beamline CMCF-BM for supporting remote diffraction data collection.

## Author contribution

A.S. and N.Z. initiated and designed the research project, performed research, analyzed the data and wrote the manuscript.

