## Supplementary information for "Structural and mechanistic basis of ubiquitous bacterial kinase signaling identifies PorX as a noncanonical substrate in *Porphyromonas gingivalis*"

**<u>Table S1</u>** - Primers used in this study.

| **Primer** | **Sequence (5' --> 3')** | **Purpose** |
|---|---|---|
| pET28 linear fw | GGATCCGAATTCGAGCTCCGTCGACAAGCTTGCGGCCGCACTCGAGCACC | pET28a-*porX*$_{PG}$ and pET28a-*ubk1*$_{PG}$ |
| pET28 linear rev | ATGGCTGCCGCGCGGCACCAGGCCGCTGCTGTGATGATGATGGTGATGATG | pET28a-*ubk1*$_{PG}$ |
| *ubk1*$_{PG}$ pET28 fw | GGCCTGGTGCCGCGCGGCAGCCATATGAATACGATCACTATCGATAGCAC | pET28a-*ubk1*$_{PG}$ |
| *ubk1*$_{PG}$ pET28 rev | GTCGACGGAGCTCGAATTCGGATCCTTAGAATGTAAGCCTGCGCTTGCCG | pET28a-*ubk1*$_{PG}$ |
| *ubk1*$_{PG}$ K39A/T40V fw | TGCTCCGATGGGTACGGGCGCGGTCACTTTCATCAAAGCCGTATGCGAGG | pET28a-*ubk1*$_{PG}$ K39A/T40V |
| *ubk1*$_{PG}$ K39A/T40V rev | CCTCGCATACGGCTTTGATGAAAGTGACCGCGCCCGTACCCATCGGAGCA | pET28a-*ubk1*$_{PG}$ K39A/T40V |
| *ubk1*$_{PG}$ D80Q fw | GACGGGCGAACTGATCTATCACTTCCAGTGCTACCGGCTCAACAAGATAG | pET28a-*ubk1*$_{PG}$ D80Q |
| *ubk1*$_{PG}$ D80Q rev | CTATCTTGTTGAGCCGGTAGCACTGGAAGTGATAGATCAGTTCGCCCGTC | pET28a-*ubk1*$_{PG}$ D80Q |
| *ubk1*$_{PG}$ E108N fw | CGATAGCGGTAGTCTCTGCTTTATCAACTGGCCGGAGCTTCTGGAGCCG | pET28a-*ubk1*$_{PG}$ E108N |
| *ubk1*$_{PG}$ E108N rev | CGGCTCCAGAAGCTCCGGCCAGTTGATAAAGCAGAGACTACCGCTATCG | pET28a-*ubk1*$_{PG}$ E108N |
| *porX*$_{PG}$ TEV p28 fw | GAAAACCTGTATTTTCAGGGCATGGAAAAAAACATGAGACCGTATACCGTAC | pET28a-*porX*$_{PG}$ |
| *porX*$_{PG}$ endrev pET28 | CTTGTCGACGGAGCTCGAATTCGGATCCTTACTTGGGTTGCATCGTAATTACGGGC | pET28a-*porX*$_{PG}$ |
| *porX*$_{PG}$ Y79A fw | GCAGAAGATCAAAGAGCTGAAACCCGCCACACCCGTCGTCATGATAACG | pET28a-*porX*$_{PG}$ Y79A, pT-groES-*porX* Y79A |
| *porX*$_{PG}$ Y79A rev | CGTTATCATGACGACGGGTGTGGCGGGTTTCAGCTCTTTGATCTTCTGC | pET28a-*porX*$_{PG}$ Y79A, pT-groES-*porX* Y79A |
| TEV p28 linear rev | GCCCTGAAAATACAGGTTTTCGTGATGATGATGGTGATGATGGTGATGATG | pET28a-*porX*$_{PG}$ |
| pET28 Kan fw | CCCATTTATACCCATATAAATCAGCATCCATGTTGGAATTTAATCGCGGC | *porX*$_{PG}$ and *ubk1*$_{PG}$ mutant construction by Gibson assembly |
| pET28 Kan rev | GCCGCGATTAAATTCCAACATGGATGCTGATTTATATGGGTATAAATGGG | *porX*$_{PG}$ and *ubk1*$_{PG}$ mutant construction by Gibson assembly |

| | | |
|---|---|---|
| Ubk1::Erm_F1_fw | CCTGGTGCCGCGCGGCAGCCATATGACCGTCTCTTTGCCCGATTCGTCAC | *Δubk1*::Erm W50 |
| Ubk1::Erm_F1_rev | GAACGGGCAATTTCTTTTTGTCATGTATGAAGTATTTACTTGGGTTGCATC | *Δubk1*::Erm W50 |
| Ubk1::Erm_Erm_fw | GATGCAACCCAAGTAAATACTTCATACATGACAAAAAAGAAATTGCCCGTTC | *Δubk1*::Erm W50 |
| Ubk1::Erm_Erm_rev | GAATGACGAACGACATCGGAGCAGGCATTACGAAGGATGAAATTTTTCAG | *Δubk1*::Erm W50 |
| Ubk1::Erm_F2_fw | CTGAAAAATTTCATCCTTCGTAATGCCTGCTCCGATGTCGTTCGTCATTC | *Δubk1*::Erm W50 |
| Ubk::Erm_F2_rev | GTCGACGGAGCTCGAATTCGGATCCTTGTCAGATGTCTGCTATTCATATTG | *Δubk1*::Erm W50 |

**Table S2** - Strains and plasmids used in this study.

| **Strain (relevant genotype)** | **Source or reference** |
|---|---|
| ***E. coli* strains** | |
| BL21 | Life Technologies |
| Top10 | Life Technologies |
| S17-1 | (1) |
| ***P. gingivalis* strains** | |
| W50 | ATCC |
| Δ*ubk1* W50 | This study |
| Δ*porX* | (2) |
| **Plasmids** | |
| pET28a (Kan$^R$) | Novagen |
| pT-COW (AmpR and TcR in E. coli; TcR in P. gingivalis; Mob+ Rep+ ) | (1) |
| pVA2198 (Em$^r$ and Sp$^r$ ) | (3) |
| pET28a-*ubk1$_{PG}$* | This study |
| pET28a-*ubk1$_{PG}$* K39A/T40V | This study |
| pET28a-*ubk1$_{PG}$* D80Q | This study |
| pET28a-*ubk1$_{PG}$* E108N | This study |
| pET28a-*porX$_{PG}$* | This study |
| pET28a-*porX$_{PG}$* Y79A | This study |
| pTCOW-groES-*porX* | (2) |
| pTCOW-groES-*porX* Y79A | This study |

**Table S3** - Data refinement statistics of the crystals. Statistic values present in parentheses correspond to the highest resolution shell data

| | |
|---|---|
| PDB code | 9ZW5 |
| Space group | $P2_12_12$ |
| Number of molecules per asymmetric unit | 2 |
| **Cell dimensions** | |
| a, b, c (Å) | 46.86, 47.79, 97.0 |
| α, β, γ (°) | 90.00, 90.00, 90.00 |
| Resolution (Å) | 42.19-2.60(2.72-2.60) |
| Rsym or Rmerge (%) | 10.7(67.0) |
| R meas (%) | 12.1(76.1) |
| Rpim (%) | 5.5(34.9) |
| I / σI | 9.6(2.2) |
| Completeness (%) | 98.8(99.8) |
| Redundancy | 4.2 |
| CC1/2 | 99.7(87.9) |
| Wavelength (Å) | 1.18 |
| **Refinement** | |
| Resolution (Å) | 42.23-2.60 |
| No. reflections (unique) | 7011 |
| Rwork / Rfree (%) | 18.92/25.44 |
| **No. of atoms** | |
| Protein | 2249 |
| Ligand/ion | 12 |
| Water | 18 |
| **B-factors** | |
| Protein | 58.69 |
| Ligand/ion | 59.1 |
| Water | 40.53 |
| MolProbity score (overall score) | 1.57 |
| Rotamer outliers (%) | 2.8 |
| Ramachandran favoured (%) | 98.92 |
| Ramachandran allowed (%) | 1.08 |
| Ramachandran outliers | 0 |
| **R.m.s. deviations** | |
| Bond lengths (Å) | 0.0074 |
| Bond angles (°) | 1.5279 |

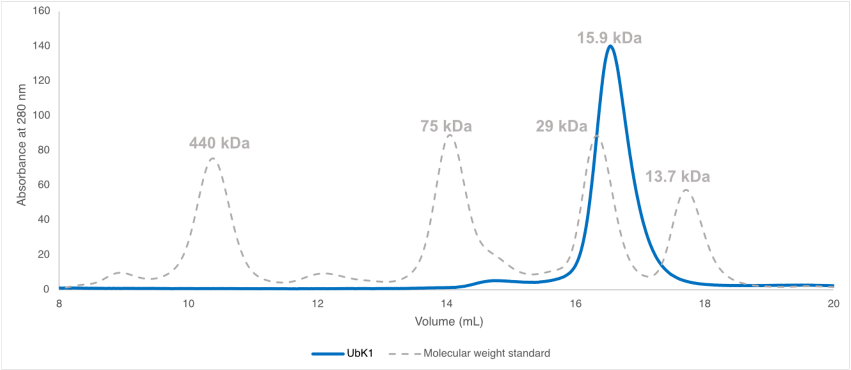


**Fig. S1** – Size exclusion HPLC of purified UbK1 compared with molecular weight standards, shows that UbK1 elutes as a monomer.

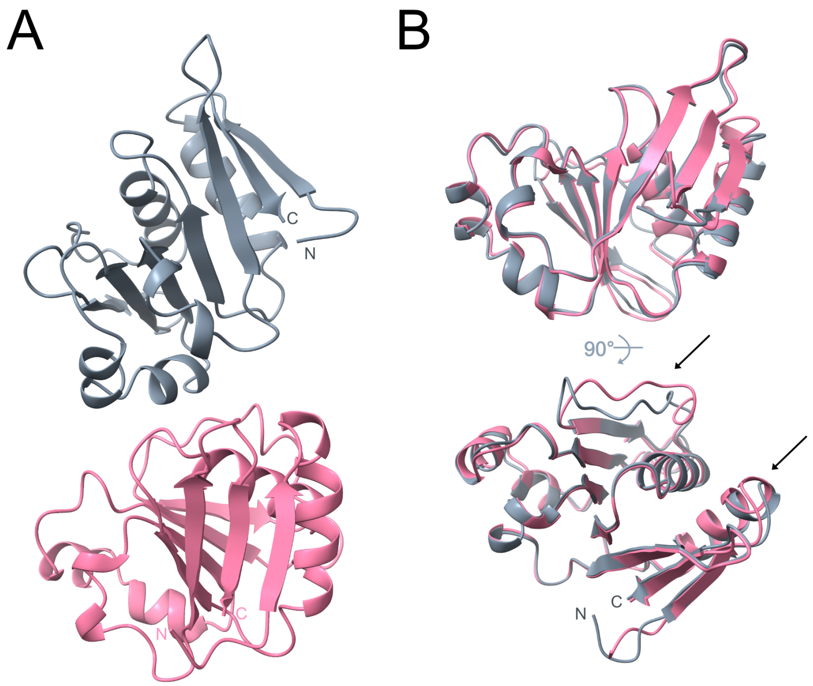


**Fig. S2** – **Crystal structure of Ubk1**. A) Asymmetric unit composition B) Superposition of the two Ubk1 monomers shows an overall highly similar architecture localized conformational heterogeneity (black arrows).

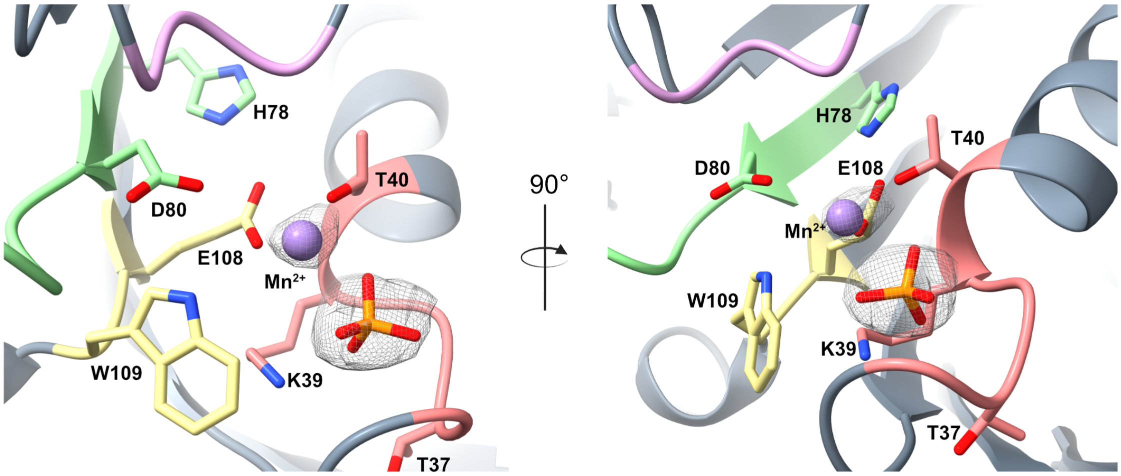


**Fig. S3 – Electron density at the UbK1 nucleotide binding site showing the bound phosphate and $Mn^{2+}$.** The motifs are coloured using the same scheme as in Fig 1. The $Mn^{2+}$ and phosphate group are shown as a purple sphere and orange sticks respectively.

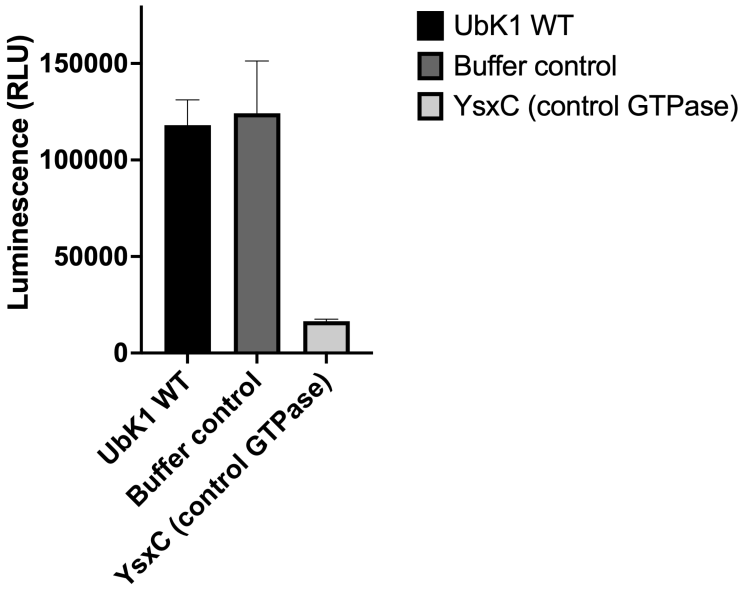


**Fig. S4** – GTPase assay showing that UbK1 does not hydrolyze GTP, exhibiting luminescence comparable to the buffer control. YsxC served as the positive control (4). In this specific assay luminescence is inversely proportional to enzyme activity.

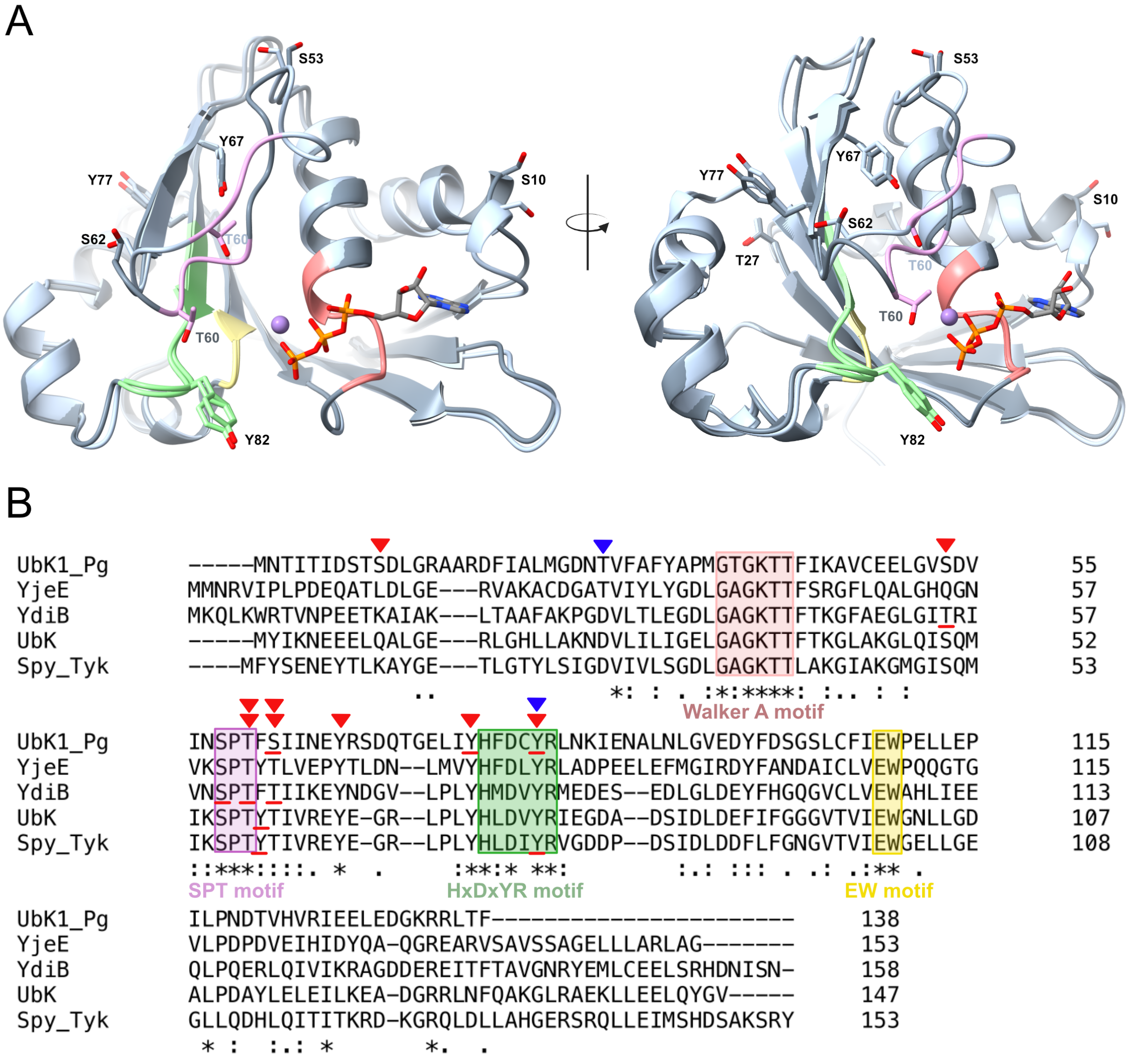


**Fig. S5** – **Mapping PorX dependent and PorX independent phosphosites on UbK1**. A) Phosphosites mapped onto an overlay of UbK1 chains (grey and light blue) docked with ATP from the *T. maritima* structure (PDB: 6N9A). ATP is shown as sticks and $Mn^{2+}$ as a purple sphere. B) Multiple sequence alignment of UbK1 with other characterized UbK-family kinases highlighting the phosphorylation landscape. Phosphosites identified in this study for UbK1 alone are marked by inverted red triangles, sites enriched or detected in the presence of PorX are indicated by inverted blue triangles, whereas previously reported sites are underlined in red. Double inverted red triangles denote the highest intensity sites detected by LCMS/MS.

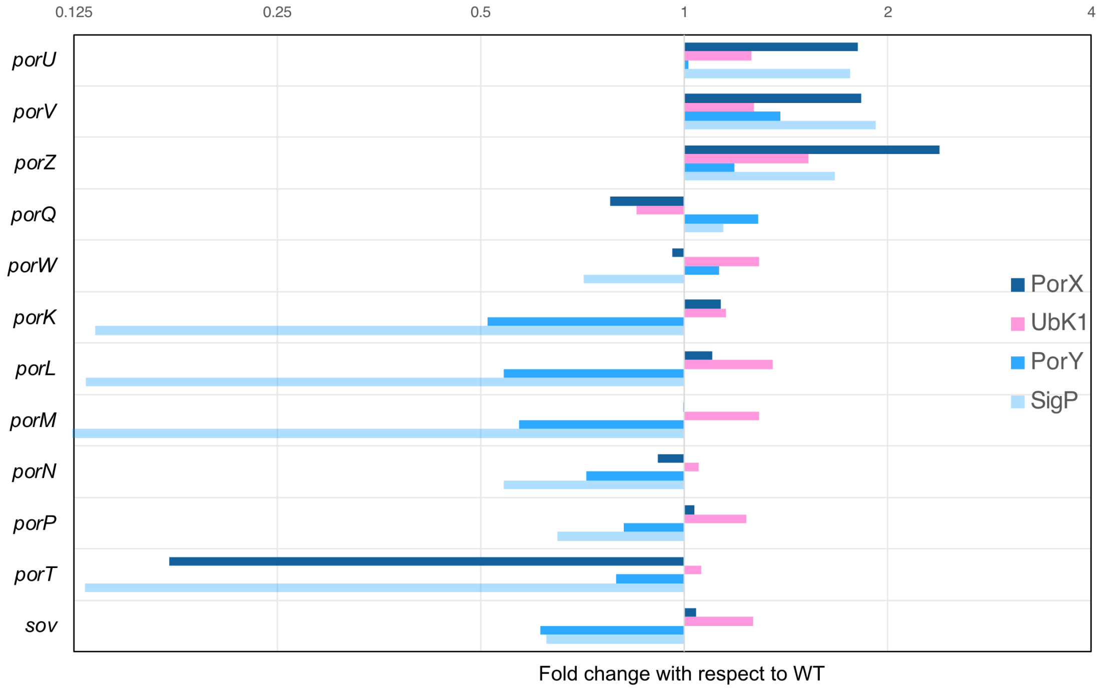


**Fig. S6** – Quantitative proteomics analysis of T9SS deletion variants shows that UbK1 abundance does not consistently correlate with T9SS gene knockouts unlike PorX and other proposed T9SS regulatory components (PorY and SigP).


1. Matsumoto-Mashimo, C., Guerout, A.-M., and Mazel, D. (2004) A new family of conditional replicating plasmids and their cognate Escherichia coli host strains. *Research in microbiology* **155**, 455-461
2. Saran, A., Kim, H.-M., Manning, I., Hancock, M. A., Schmitz, C., Madej, M., Potempa, J., Sola, M., Trempe, J.-F., Zhu, Y., Davey, M. E., and Zeytuni, N. (2024) Unveiling the molecular mechanisms of the type IX secretion system's response regulator: Structural and functional insights. *PNAS Nexus* **3**
3. Bolivar, F., and Backman, K. (1979) Plasmids of Escherichia coli as cloning vectors. in *Methods in enzymology*, Elsevier. pp 245-267
4. Ni, X., Davis, J. H., Jain, N., Razi, A., Benlekbir, S., McArthur, A. G., Rubinstein, J. L., Britton, R. A., Williamson, J. R., and Ortega, J. (2016) YphC and YsxC GTPases assist the maturation of the central protuberance, GTPase associated region and functional core of the 50S ribosomal subunit. *Nucleic Acids Res* **44**, 8442-8455
